# Self-referencing rates in biological disciplines

**DOI:** 10.1101/2023.04.19.537562

**Authors:** Sean M. Cascarina

**Affiliations:** Department of Biochemistry and Molecular Biology, Colorado State University, Fort Collins, CO 80523, USA

## Abstract

The use of citation counts (among other bibliometrics) as a facet of academic research evaluation can influence citation behavior in scientific publications. One possible unintended consequence of this bibliometric is excessive self-referencing, where an author favors referencing their own publications over related publications from different research groups. Peer reviewers are often prompted by journals to examine whether references listed in the manuscript under review are unbiased, but there is no consensus on what is considered “excessive” self-referencing. Here, self-referencing rates are estimated across multiple journals in the fields of biology, genetics, computational biology, medicine, pathology, and cell biology. Median self-referencing rates are between 8-13% across a range of journals within these subdisciplines. However, self-referencing rates vary as a function of total number of references, number of authors, author status/rank, author position, and total number publications for each author. These results may serve as useful statistical guidelines for authors, editors, reviewers, and journals when considering referencing practices for individual publications.

## Introduction

The use of bibliometrics (i.e. quantitative metrics applied to publication records) to assess research success has become commonplace. In principle, data-driven methods of evaluating individual researchers could contribute to more objective, consistent, comprehensive, and pragmatic assessments [1], but they are also prone to manipulation [2], can inadvertently distort research incentives [3], and can be misused in the decision-making process [4]. For these reasons, recent guidelines on individual researcher assessment advocate integrating multiple, complementary quantitative measures [5], as well as balancing these quantitative measures with qualitative, expert assessment of researcher contributions [3,4].

Bibliometrics are often designed to approximate the “impact” of an individual within a scientific discipline [6]. One such statistic, the number of citations to published work by a researcher, is often an integral part of bibliometrics that attempt to estimate impact [6–8]. While it is reasonable to expect influential ideas and discoveries to be cited frequently, the potential motivations for citing a particular publication are numerous and are not always based purely on the scientific content of that publication [7]. Additionally, citations can be influenced by the authors themselves via self-referencing or “citation rings” [9–13], by reviewers [14–17], by editors [18,19], and by journals [19–24] (for additional discussion on all of these categories, see [2,25]). Influence of citations is not always unethical, but requires a delicate (and often subjective) balance in appropriate cases. For example, peer reviewers can recommend adding references to their own work when evaluating a manuscript, but this should only occur in cases where the references are truly pertinent to the manuscript, and exclusion of suggested references generally should not be grounds for rejection [26]. Similarly, authors often reference their own prior publications when it is a continuation of previous lines of research (which helps readers contextualize the work), but excessive self-referencing can be considered inappropriate and is sometimes discouraged/explicitly restricted in journal guidelines or instructions to peer reviewers. Authorship and citation practices also vary by discipline (e.g. [27–29]), making it difficult to ascertain what a “normal” self-referencing rate is in a particular subdiscipline.

Here, self-referencing rates are evaluated at the per-article level for >94k publications from a subset of selected journals representing a cross-section of biological disciplines over the past 20 years. Shifts in self-referencing rate distributions are examined as a function of journal, publication date, and total number of works referenced. Factors associated with unusually high self-referencing rates such as author status and total number of publications are also explored. These observations provide a range of quantitative guidelines for authors, reviewers, editors, and journals based on recent historical self-referencing practices, while also highlighting instances of situationally appropriate extreme self-referencing.

## Results

### Self-Referencing Rates Are Similar Across Biology-Related Disciplines

PubMed is a popular literature database for the life sciences, composed of >30 million publications and typically searched >2.5 billion times per year (https://www.nlm.nih.gov/bsd/medline_pubmed_production_stats.html). Although PubMed provides a complete set of downloadable, open-access articles, publications in a small subset of journals were used instead because individual PubMed queries improve author disambiguation for purposes of this study (see Methods). Analyses were initially focused on the *PLOS* (Public Library of Science) family of journals for five main reasons: 1) the core *PLOS* journals were founded shortly after the inception of PubMed, ensuring that most publications would be available for retrieval in automated searches; 2) specialized *PLOS* journals span a variety of biology-related disciplines (i.e. general biology, genetics, pathology/pathogens, medicine, and computational biology); 3) the *PLOS* journals have been open access since their inception, potentially increasing the likelihood of retrieving a complete set of publications; 4) the *PLOS* journals strongly favor the publication of primary research; and 5) the volume of publications in the core *PLOS* journals and all referenced works in those publications was computationally feasible.

For the purposes of this work, the operational definitions of “self-referencing” and “self-citation” are adopted from Szomszor *et al*. [28]. Specifically, self-referencing refers to the inclusion of one’s own prior publication in the reference list of a later publication. The per-article “self-referencing rate” can be estimated for each author individually and is defined as the percentage of references in each publication that were also (co-)authored by that individual. To examine self-referencing rates, an author list was retrieved for each publication in the core *PLOS* journals (“primary publication”) and cross-referenced against the author lists for each publication cited (“referenced publication”) in that article. The self-referencing rate assigned to each primary publication was then simplified as the maximum self-referencing rate among authors of that primary publication, which was typically achieved by the last/corresponding author of each publication (evaluated further in a later section). Estimated self-referencing rates are provided in Table S1. Computational estimates of self-referencing rates are extremely consistent with manual calculations performed for a randomly selected subset of publications (*R^2^* = 0.976; Fig S1 and Table S2).

Self-referencing rate distributions are similar across the *PLOS* journals (Fig 1A and Table S3), with median self-referencing rates between 8.2-11.4% and long, positively skewed tails. *PLOS Pathology* and *PLOS Medicine* exhibit the greatest positive skew and average self-referencing percentage (13.4% and 13.5%, respectively). *PLOS Computational Biology* exhibits both the least positive skew and lowest mean self-referencing percentage (10.1%). To determine whether self-referencing rates are changing over time, publications were stratified by journal and publication year. No clear trend in median self-referencing rate is apparent for any of the *PLOS* journals (Fig 1B and Table S3), suggesting that self-referencing rates have been relatively stable over the lifetime of these journals.

**Figure 1.**
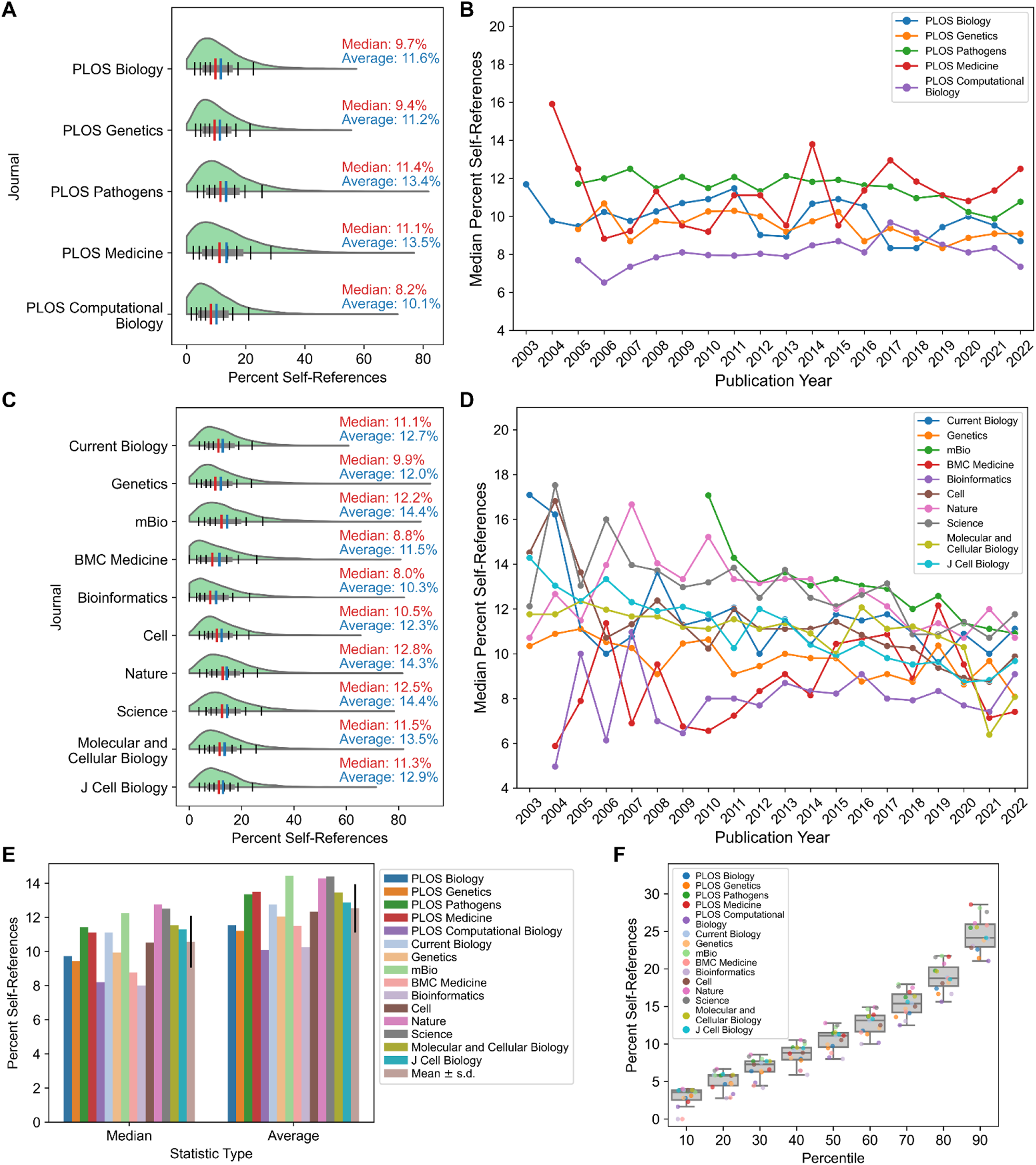
Self-referencing rate is stable across time and journal. (A) Self-referencing rate distributions for publications in core *PLOS* journals. (B) Year-stratified median self-referencing rates since 2003 for the core *PLOS* journals. (C) Self-referencing rate distributions for non-*PLOS* journals. (D) Year-stratified median self-referencing rates since 2003 for non-*PLOS* journals. (E) Comparison of median self-referencing rates and average median self-referencing rates (±s.d.) across all journals evaluated. (F) Distributions of percent self-referencing rates at 10-percentile intervals for all journals evaluated. To mitigate high self-referencing rates for publications with few total references, only publications with 20 or more total references were included in the analyses for all figure panels. For panels E and F, *PLOS*-matched journals are indicated in identical colors with lighter hues.

While the *PLOS* journals provided a suitable case study for the reasons outlined above, it is possible that publisher practices or policies could influence self-referencing rates during the peer review or editorial stages. For each specialized *PLOS* journal, a non-*PLOS* journal with similar subdiscipline, impact factor, and number of publications in the same 20-year timeframe was selected for comparison (Fig S2A,B). Two additional cell biology journals were included to account for this large life science subdiscipline that does not have a dedicated *PLOS* journal. Furthermore, the journals *Cell*, *Nature*, and *Science* were included to explore potential effects correlated with high impact factors. Self-referencing distributions for the non-*PLOS* journals are largely similar to those of *PLOS* journals (Fig 1C and Table S3), indicating that the publisher does not strongly influence self-referencing rates within subdisciplines, though important minor differences are observed. Like its corresponding *PLOS* journal, *Bioinformatics* has the lowest median self-referencing rate and less positive skew compared to most other journals. However, in contrast to its *PLOS* counterpart, BMC Medicine has a low median self-referencing rate and less positive skew compared to many journals, which may in part be attributed to a higher number of total references compared to *PLOS Medicine* (Fig S2C). *Nature* and *Science* have the highest median self-referencing rate and greatest positive skew of all journals, potentially due to publication of research in non-life-science disciplines, which can have substantially larger author lists and higher self-referencing rates (e.g. physics; [10]). Median self-referencing rates for these journals span a broader range in the early 2000’s but exhibit some degree of convergence into a smaller range in recent years (Fig 1D and Table S3), which may suggest that publisher practices and/or publishing conventions are evolving over time and becoming more standardized.

Importantly, cumulative comparison of self-referencing rates across all journals evaluated indicates that median self-referencing rates fall within a narrow range (Fig 1E) with similar distributions (Fig 1F) in biology-related disciplines, though there are minor differences between fields. These estimates are largely consistent with average self-referencing rates in neurology, neuroscience, and psychiatry reported in a recent preliminary study [30], albeit with mean self-referencing rates slightly higher in neurology and psychiatry compared to all journals evaluated here.

### Self-Referencing Rates Are Dependent on the Total Number of References

Publications in *PLOS Medicine*, *Nature*, and *Science* have some of the highest median and average self-referencing rates (Fig 1). However, these publications also tend to have notably fewer total references, on average, compared to publications in the other journals evaluated (Fig S2C). Higher self-referencing rates might be expected for publications with fewer total references since, in those instances, relatively few citations to prior work could constitute a high percentage of total references.

To examine this directly, trends in the median (Q2) and interquartile range (Q1 and Q3) were studied as a function of the total number of references. *A priori*, quartile values were expected to decrease for publications with a larger number of total references since high percent self-reference values would be more difficult to achieve as total references increase. Indeed, for all journals, quartile values typically decrease steadily as the total number of references increases (Fig 2 and Table S4), indicating that the distributions of self-referencing rates shift toward lower percent self-reference values. Total references have a relatively strong effect on self-referencing rates, with most journals exhibiting a 5 to 10 percentage-point decrease in median self-referencing rate as total references increase. However, not all journals exhibit identical behavior. For example, *PLOS Biology* starts with a relatively low median self-referencing rate (∼11.9%) but is less sensitive to total references than most journals, with the lowest self-referencing rate reaching ∼6.5% and corresponding a range of only ∼5.4% (the second-lowest range of all journals). *Molecular and Cellular Biology* (*MCB*) starts with the highest median self-referencing rate (∼17.1%) for publications with a low number of total references but exhibits a steep decline, eventually resulting in one of the lowest self-referencing rates (∼5.5%; range ∼11.6%) for publications with a high number of total references (Fig 2). Similarly, *PLOS Medicine* exhibits a steep decline in self-referencing rates as total references increase: however, a large number of *PLOS Medicine* publications have relatively few references (Fig S2C), suggesting that its high overall median self-referencing rate is driven predominantly by the practice of citing fewer publications. In contrast, publications in *Nature* and *Science* simultaneously tend to have few references (Fig S2C), high initial self-referencing rates at a low number of total references (Fig 2), and early plateaus (and sometimes even an increase) in quartile values as a function of total references; all three factors likely contribute to their unusually high median self-referencing rates. Publications in *Cell* also exhibit relatively high initial self-referencing rates at a low number of total references (Fig 2), but the majority of publications have a larger number of total references (Fig S2C). Consequently, the majority of *Cell* publications are found within a region of the curve with a lower self-referencing rate, driving down the overall mean and median self-referencing rates relative to *Nature* and *Science* (Fig 1C,E).

**Figure 2.**
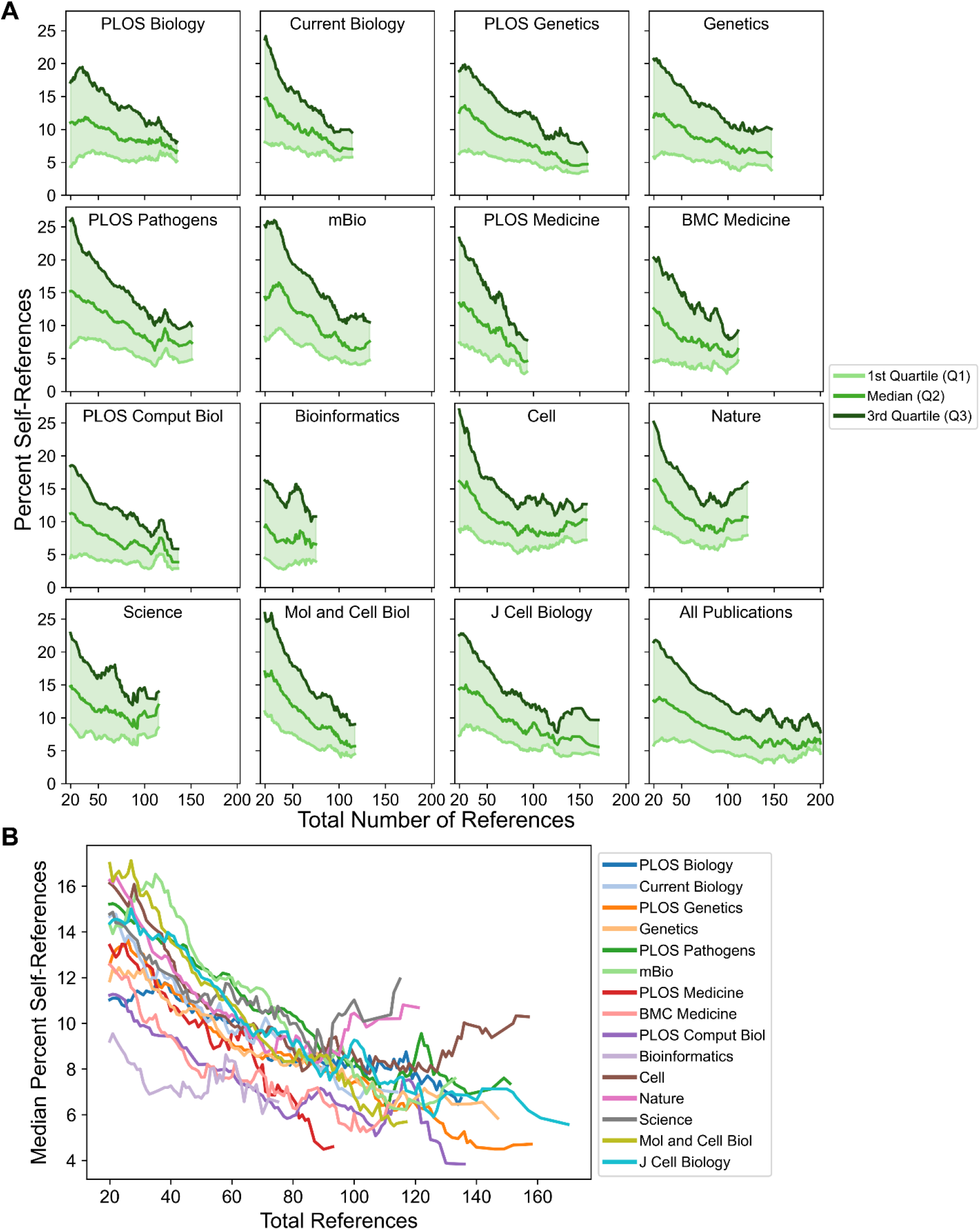
Changes in self-referencing rate distributions as a function of total references. (A) Distributions of percent self-references (as indicated by values corresponding to the Q1, median, and Q3) for each journal shift as a function of total references. (B) Direct comparison of the median percent self-reference value across journals as a function of total references. For all analyses, curves were smoothed by calculating a sliding mean with a window size of 11 (see Methods for additional details).

In summary, these observations suggest that self-referencing rates across many journals are strongly dependent on the total number of references. The degree and shape of the dependence differ by journal and are likely influenced by journal policies related to total allowable references or total article length restrictions, among other factors (see Discussion). Therefore, variation in total references results from both individual author habits and systemic factors that affect total references.

### Total Number of Authors has Only A Minor Effect on Maximum Self-referencing Rates

Differences in the total number of authors on each publication are observed across journals (Fig S2D). Previous bibliometric studies indicate a correlation between the total number of authors on a publication and the self-referencing/self-citation rate in some fields [29,31,32]. However, these studies use a different metric: the *collective* self-referencing rate, which is defined as the percentage of references for which any author on the primary publication was also an author of the referenced publication. This is in contrast to the *individual* self-referencing rate (e.g. [11,27,28]), which is calculated independently for each author and is here used to determine the maximum individual self-referencing rate. Compared to the maximum individual self-referencing rate, the collective self-referencing rate is likely more strongly correlated with the total number of authors. Nevertheless, the total number of authors could still affect the maximum individual self-referencing rate by increasing the probability that a higher maximum self-referencing rate is achieved, since each additional author may have unique self-referencing habits and a unique pool of prior publications that could be referenced.

Self-referencing rates exhibit a weaker dependence on the total number of authors and more-cross journal variability (Fig 3 and Table S5) when compared to the dependence on the total number of references. Nearly all journals exhibit at least a slight increase in typical self-referencing rates as the number of authors increases. However, for many journals such as *PLOS Biology, Current Biology, mBio, PLOS Computational Biology, Cell, MCB,* and *J Cell Biology* this increase does not exceed more than ∼3 total percentage points gained. For the journals *PLOS Genetics, Genetics, PLOS Pathogens, Bioinformatics,* and *Nature*, moderate increases of ∼3-5 percentage points are observed. Notably, both medical journals (*PLOS Medicine* and *BMC Medicine*) exhibit large increases of ∼7-10 percentage points as the total number of authors increases, potentially indicating a domain-specific practice. Publications in *Science* also exhibit a large increase in self-referencing rate, but only starting at publications with ∼30 or more authors.

**Figure 3.**
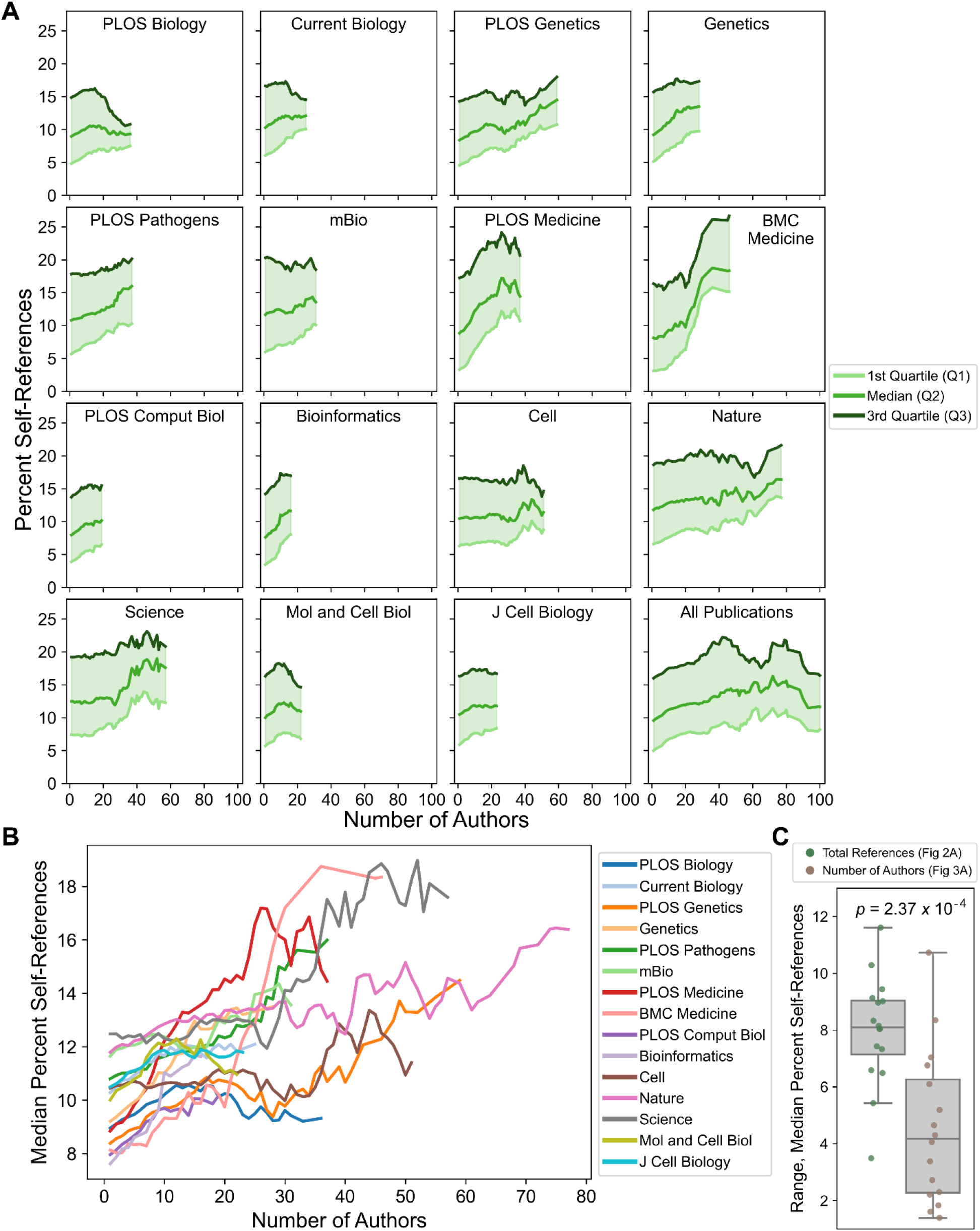
Changes in self-referencing rate distributions as a function of total number of authors listed on each publication. (A) Self-referencing rate distributions (Q1, median, and Q3) for each journal shift as a function of total number of authors. (B) Direct comparison of the median percent self-reference value across journals by total number of authors. For all analyses, curves were smoothed by calculating a sliding mean with a window size of 11 (see Methods for additional details). (C) Distributions of the statistical range for median self-referencing rates. Each dot represents the self-referencing rate range for a single journal as a function of total references (green) or number of authors (brown). The indicated *p*-value was calculated from a two-sided *t*-test.

One possible explanation for the weak relationship between percent self-references and the total number of authors is that authors are unlikely to contribute equally to the self-referencing rate. For example, “senior authors” (i.e. principle investigators or lab directors) are presumably the most likely to achieve the highest self-referencing rate since they typically have the longest history of closely related publications, but senior authors usually constitute a small fraction of the author list. In contrast, “junior authors” (e.g. undergraduates, graduate students, post-docs, and research faculty) often have fewer related publications compared to the senior author but constitute the majority of the author list. From this perspective, self-referencing rates would be most strongly influenced by the addition of senior authors, as occurs for collaborative publications. Therefore, the total number of authors is likely an imperfect proxy for the number of senior authors included in the collaboration.

To examine this more directly, publications were binned according to the number of authors. In a conventional research group architecture, at least one junior scientist participates in the research project for every senior scientist involved. Therefore, publications with three or fewer authors were generally assumed to originate from a single laboratory and have a single senior author. Indeed, the maximum self-referencing rate is achieved by the last author for ∼85% of publications with three or fewer total authors (Fig 4A), suggesting both that the large majority of publications in this group likely originate from a single laboratory and that the senior author is nearly always the highest self-referencing author. In the remaining 15% of publications, the last author was sometimes not the corresponding author, suggesting that the senior author is still the maximum self-referencing author in a portion of these instances. Publications with four or more total authors are considered potentially collaborative, with the probability of collaboration expected to increase as the total number of authors increases. For all total-author bins examined, the last author achieves the maximum self-referencing rate for most of the publications. The share of publications progressively decreases for all total-author bins as a function of author position when moving inward from the last author, confirming that the most senior authors are indeed the most likely to achieve the highest self-referencing rate, and that less senior authors are progressively less likely to achieve the maximum self-referencing rate.

**Figure 4.**
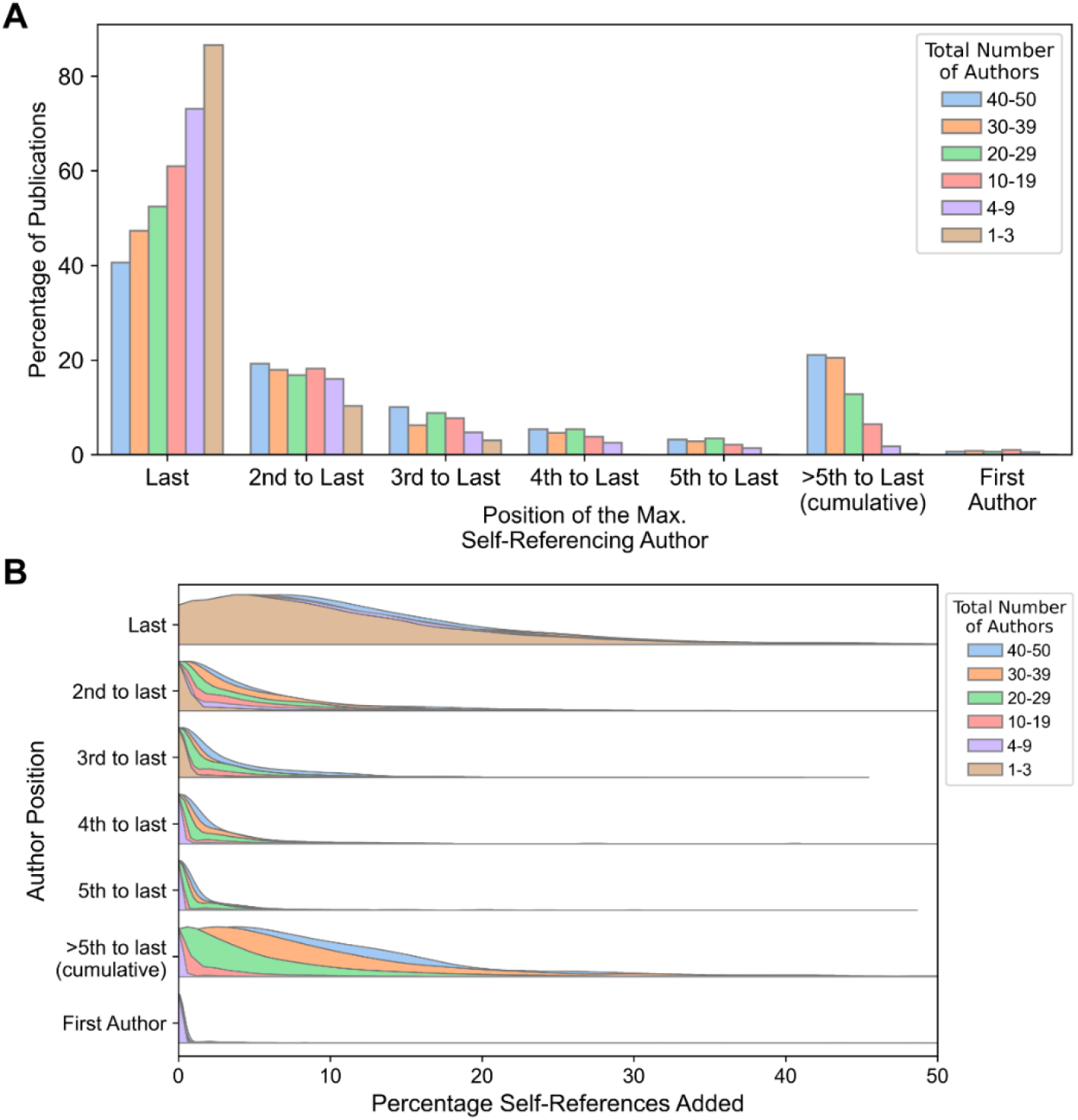
Self-referencing statistics by author list position. (A) The share of publications (as a percentage) for which the maximum self-referencing rate was assigned to each author for the final five authors and the first author, stratified into bins by total number of authors. (B) Distributions of the percentage of self-references added to the collective self-referencing rate by each author position. The percentage of self-references added is calculated sequentially by determining the individual self-referencing rate for each author while excluding self-referenced publications that were already accounted for in a preceding author, starting with the last author and moving inward (i.e. last author, 2^nd^ to last author, 3^rd^ to last author, etc.). For both panels, the author on single-author publications is classified as the last author. In cases of ties between authors, the maximum self-referencing rate was attributed to the most senior author (i.e. the author nearest to the end of the author list), as this reflects the number of unique references contributed by each additional author.

Some senior authors frequently collaborate with the same senior co-author(s), which could lead to situations where the sets of self-references are highly overlapping. In these instances, even a single additional self-reference could shift the maximum self-referencing rate away from the last author to another senior author. For example, although the last author may not achieve the maximum self-referencing rate among collaborative publications (e.g. 30-39 authors), they may still be responsible for the majority of the self-references, and the share of unique additional self-references associated with non-last authors may be minimal.

For each publication, percent self-references were sequentially calculated starting with the last author. For each subsequent author (moving inward from the last author), a self-reference was only included in the calculation if it had not already been included as a self-reference for the preceding authors, effectively exploring the unique contribution of each author to a collective self-referencing rate. The last author tends to contribute the overwhelming majority of self-references for all total-author bins (Fig 4B and Table S6). Surprisingly, this is largely independent of the total number of authors, with only a marginal rightward shift in the percentage of self-references distribution for publications with 40-50 authors. This suggests that self-referencing habits of the most senior author tend to be insensitive to the number of additional co-authors (including senior authors), and may reflect a certain baseline self-referencing that authors deem appropriate for describing prior work. However, for subsequent authors (i.e. 2^nd^ to last, 3^rd^ to last, etc.), the unique contribution to percentage of self-references is dependent on both the total number of authors and their proximity to the end of the author list.

Collectively, these results corroborate prior observations that self-referencing is dependent on the total number of authors. However, the majority of unique self-references for most publications are attributable to the last author in a manner that is insensitive to the total number of authors (with the possible exception of certain subdisciplines such as medicine). Minor additional self-references are subsequently contributed by senior co-authors, and this contribution scales with the total number of authors and proximity to the end of the author list.

### Self-referencing Rates Are Related to Author Rank and Total Number of Publications

One possible explanation for high self-referencing rates is an unusually significant contribution to a particular field. While “contribution significance” is at least partially subjective and difficult to measure, two important components of contribution significance are breadth (e.g. the total number of publications associated with an individual) and depth (e.g. the degree of impact, on average, associated with an individual) [6]. A recently published database of highly ranked researchers [9,33] contained both the total number of publications by each listed researcher and a composite score (based on six distinct citation measures; [5]) used to rank each researcher, which are suitable estimates of breadth and depth.

For all authors present in the ranked-authors database, the relationship between self-referencing rates for the maximum self-referencing author and the total number of publications for that author was explored. Very high self-referencing rates (>30%) typically occur at the low end of the total-publications spectrum, with a dense cluster between ∼50-250 total publications (Fig S3A,B). However, a notable portion of the scatterplot extends toward higher total publications (Fig S3A). To better understand how the distribution of self-referencing rates changes as a function of total publications by an author, quartile values were calculated for self-referencing rate distributions as a function of total publications. Quartile values for self-referencing rate distributions rise sharply until ∼200 total publications (Fig 5A). Beyond 200 publications, quartile values continue to rise on average, but at a much slower rate. This is consistent with previous observations [34] that higher self-referencing rates tend to occur for authors with more total publications, but suggests a more nuanced relationship between total publications and self-referencing rates: beyond a certain point, the effect of each additional publication on self-referencing rate diminishes.

**Figure 5.**
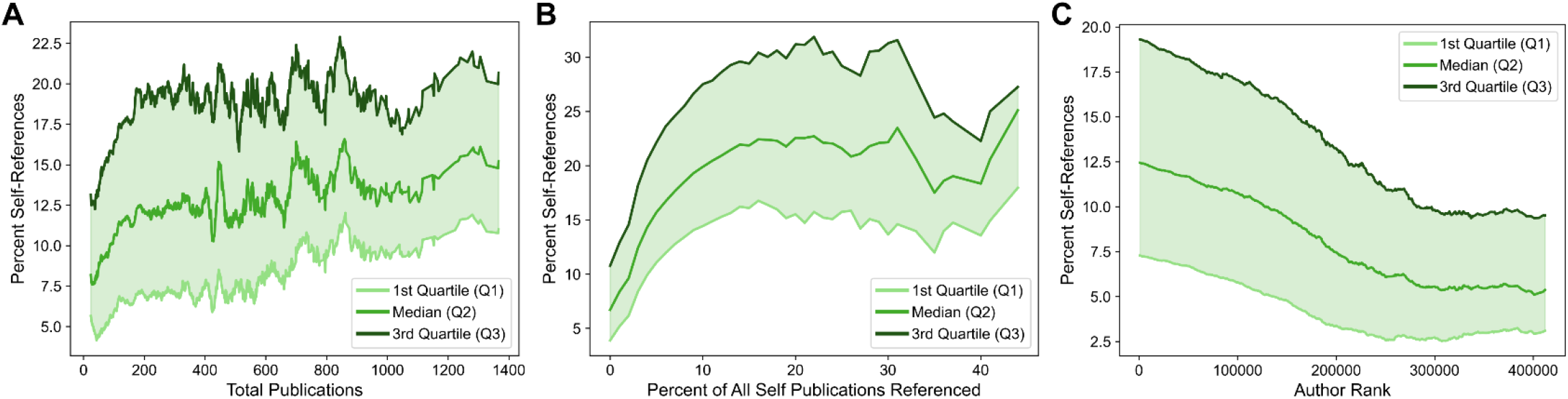
Self-referencing rates among ranked authors as a function of total publications, percentage of self publications referenced, and author rank. Self-referencing rate quartile values (Q1, median, and Q3) shift as a function of total publications (A), percentage of all “self” publications referenced (i.e. those published by the author being evaluated) (B), and author rank (C). For all panels, curves were smoothed by sliding means (see Methods). Only authors appearing in the ranked-authors database [9,33] are included in the analyses.

Hypothetically, authors could attempt to reference most or all of their prior work in their own publications, which would likely result in both a high self-referencing rate and a high percentage of the author’s total existing publications referenced. The self-referencing rate quartile values exhibit an initial increase with an increasing percentage of total publications referenced but plateaus at ∼15% of total publications referenced (Fig 5B), at which point the median self-referencing rate stabilizes at ∼22-23%. The magnitude of the increase in self-referencing rate is larger than that observed as a function of total publications. However, instances with both a high self-referencing rate and a high percentage of total publications referenced represent a small minority of publications (Fig S3C,D), suggesting that most authors are not attempting to cite a large proportion of their work at once. Since a simultaneously high self-referencing rate and high percentage of publications referenced would be difficult to achieve for authors with a large number of total publications, this suggests that high self-referencing rates may also be achieved by citing a larger fraction of prior work, particularly for authors with fewer total publications.

The ranked-authors database includes a ranking that is independent of self-citation. This aids in mitigating downstream effects of authors that boost their apparent impact by heavily referencing their own work (though non-self-references are not entirely independent of self-references; [11,35]). A steady shift from higher to lower self-referencing rates is observed when traversing from high-ranking authors to lower-ranking authors (Fig 5C). Very high self-referencing rates are also preferentially clustered toward authors with the highest rank (Fig S3E,F). It is worth noting that high-ranking authors also tend to have a larger number of total publications and that a higher number of total publications is likely to garner more total citations (Fig S4A-C). However, the decrease in self-referencing rate as author rank decreases is still observed when authors are stratified based on the total number of publications, although the magnitude of the effect progressively decreases for author groups with larger total publications (Fig S4D-F). This suggests that the correlation between self-referencing rate and author rank is at least partially independent of the total number of publications. Therefore, high self-referencing rates often correspond to works from unusually influential authors, as judged both by total publications and non-self, citation-based author rank.

Although clear trends are observed within the ranked-authors database, these authors still constitute a biased subset of researchers, since all authors included in the database are considered highly influential. Therefore, self-referencing rates for authors included in the ranked-authors database were compared to self-referencing rates for authors excluded from the ranked-authors database. The distribution of self-referencing rates is significantly shifted toward higher self-referencing percentages for ranked authors compared to non-ranked authors (Fig 6A). Additionally, the percentage of publications for which the maximum self-referencing author appears in the ranked-authors database steadily increases among publications with increasingly high self-referencing rates (Fig 6B). For example, only ∼10% of publications with 0% self-references correspond to publications where the maximum self-referencing author appears in the ranked author database. However, this number jumps to nearly 60% among publications with ∼20% self-references, and further increases to ∼70-80% for publications with ≥40% self-references. Thus, highly influential authors, regardless of their rank in the database, exhibit higher self-referencing rates than non-ranked authors

**Figure 6.**
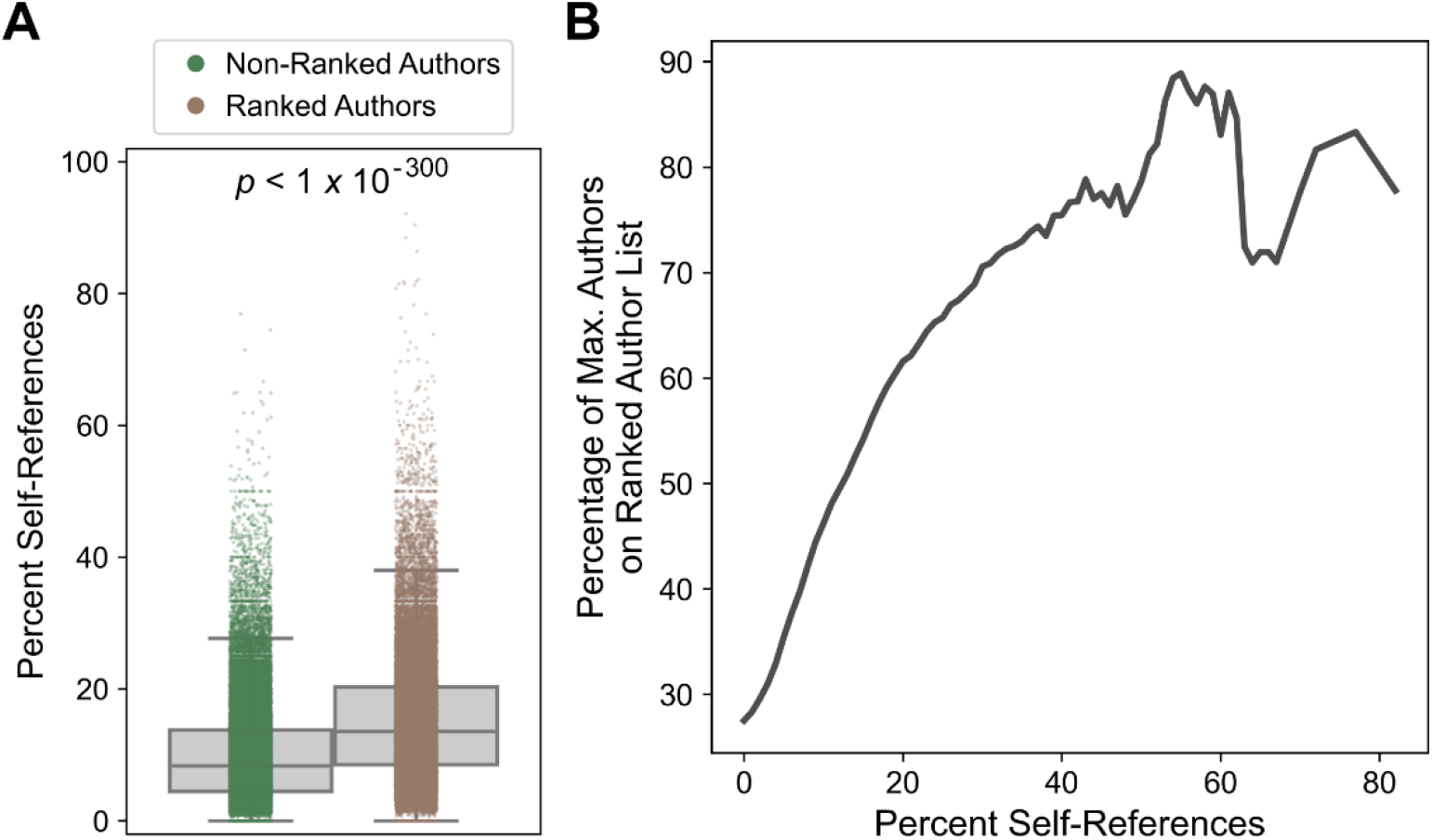
Comparison of self-referencing rates for ranked and non-ranked authors. (A) Self-referencing rate distributions for ranked and non-ranked authors. The indicated *p*-value was calculated from a two-sided *t*-test and was below the precision limit of the software. In cases of ties, the most senior author involved in the tie was evaluated for inclusion/exclusion in the ranked-authors database. (B) Percentage of the maximum self-referencing authors that appear on the ranked-author list as a function of minimum percent self-references.

## Discussion

### Self-Referencing in Biology-Related Disciplines

Self-referencing in science often serves a legitimate purpose: since most scientific endeavors build upon previous work from the same research group, appropriate self-referencing can help readers contextualize a publication by highlighting prior foundational work [28,36,37]. From this perspective, strict, one-size-fits-all rules limiting self-referencing could lead to referencing practices that misrepresent existing literature. Based on a limited poll, most researchers are against journals “policing” self-referencing [10]. On the other hand, the use of citations as a simple summary statistic influencing professional outcomes (e.g. grant funding, hiring/promotion decisions, recognition, etc.) incentivizes researchers to disproportionately cite their own work, which also misrepresents existing literature [28,36]. Many of the cited studies acknowledge the potential existence of “excessive” self-referencing, though opinions on precise definitions and criteria for “excessive” vary. Therefore, self-referencing practices require a delicate balance that must be navigated by research authors and journals alike.

The statistical analyses presented here of self-referencing rates over the past 20 years reveal modern, unwritten conventions among researchers in biology-related fields, and how these conventions continue to evolve. In principle, self-referencing rates could be automatically detected by journals upon manuscript submission using computational methods similar to the approach utilized here, and estimated percentile values for self-referencing rates could be provided to peer reviewers to inform their assessment. In that sense, journals would not be “policing” self-referencing: rather, they would be providing contextual information to the peer review experts primarily responsible for evaluating a manuscript – a task that they are often prompted to do anyway, without this contextual information. Even in the absence of an automated system, peer reviewers can estimate the self-referencing rate for each manuscript and compare it to these statistical benchmarks. Importantly, self-referencing rates and their corresponding percentile benchmarks could be normalized by the total number of references: this would prevent authors from artificially lowering their percentage of self-references by including extraneous, non-self-references to inflate the total number of references. Journals, reviewers, and authors could also consider the total number of publications by the author(s), which could justifiably increase self-referencing rates if the authors have a relatively large number of citable works; but even this should be considered thoughtfully as it can be gamed by authors [2].

Although these statistical benchmarks are indicative of modern self-referencing conventions, they do not preclude the possibility of publications with exceptionally high self-referencing rates. Such situations could include researchers working on emerging or underexplored topics, researchers with exceptional influence in a particular field (e.g. “founders” of a field of study), or publications with relatively few references, which may be short, narrowly focused publications. For example, two publications were identified with an estimated self-referencing rate >90%: however, upon manual inspection, both publications appear to be invited papers accompanying an award to an individual researcher and were intended to be an autobiographical account of their discoveries in a particular field. Even with these exceptions, self-referencing rates rarely exceed ∼20% (surpassed only by the 90^th^ percentile across journals in Fig 1F), indicating that situationally high self-referencing rates are uncommon.

To the best of my knowledge, none of the journals evaluated provide concrete guidelines or restrictions on self-referencing which, again, appears to actually be preferred among researchers [10]. A lack of explicit self-referencing policies is also advantageous for the present study, since *a priori* restrictions to self-referencing could affect the observed distributions. However, some journals have policies that indirectly affect self-referencing rates. For example, *Nature* and *Science* recommend 50 or fewer total references for their main article types which, in light of the relationship between total references and self-referencing rates (Fig 2), may also partially explain the higher mean and median self-referencing rates and more positive skew in the distribution of self-referencing rates. *Bioinformatics* does not explicitly limit the total number of references but imposes length restrictions that include the references, which indirectly affects self-referencing rates. *Cell*, *J Cell Biology*, *MCB*, *mBio* and *Current Biology* impose article length restrictions but exclude the references from these restrictions. The remaining journals (*PLOS* journals, *Genetics,* and *BMC Medicine*) do not restrict article length or references. The only journal that explicitly addressed self-referencing was *BMC Medicine*, whose editorial policies included a statement that excessive self-referencing and citation manipulation are strongly discouraged and grounds for rejection, but they do not define what constitutes “excessive” self-referencing. This lone example amounts to only ∼6.7% of the journals evaluated here having an explicit self-referencing policy, which is lower than estimates in other fields such as critical-care medicine [38] and anesthesiology [39], albeit with a relatively small sample size in the present study.

Some journals explicitly restrict self-referencing to a certain percentage, although this does not currently seem to be a common practice. For example, in their author guidelines *Current Opinion in Cell Biology* expresses a hard upper limit of 20% self-references, while *Langmuir* (a chemistry journal) states that a 25% self-reference upper limit is imposed but that exceptions can be made by consulting with the editors (both sets of guidelines accessed on 1/31/2023). Neither journal specifies whether these upper limits apply to *individual* or *collective* self-referencing rates. Assuming these limits refer to maximum individual self-referencing rates, they would roughly correspond to the 85^th^-95^th^ percentile for the biological journals evaluated in this study. If the limits correspond instead to collective self-referencing rates they are potentially more restrictive, particularly for studies with many authors or multiple senior authors. Regardless, such limits – based on self-reference rate cutoffs rather than percentiles – have the potential to change self-referencing distributions over time, eventually resulting in new “normal” ranges. Furthermore, these limits do not account for the relationship between self-referencing percentages and the total number of references in the manuscript (Fig 2), the total number of authors on the publication (Figs 3,4), or the total number of citable publications associated with the self-referencing author (Fig 5). This effectively penalizes publications with shorter reference lists, more total authors, or a large number of citable prior publications, respectively. These factors, in conjunction with statistical benchmarks, enable a more informed and contextualized assessment of self-referencing rates by authors, peer reviewers, editors, and journals.

Finally, I wish to make clear that I am not advocating for or against any particular policy regarding self-referencing: these data are provided merely to aid in the development of informed decisions by relevant experts, which will depend on each situation. Discussion of individual researchers was also intentionally omitted to avoid shaming or “criminalizing” [10] self-referencing, and because a focus on specific individuals is here considered less important and substantive than the broader statistics of self-referencing.

### Limitations of the Study

There are a number of limitations worth discussing. First, although failed PubMed queries were repeated multiple times (see Methods section), a small subset of PubMed queries for primary publications and referenced publications were still not successful. Failed queries most often occurred for non-research article types (e.g. editorials, author corrections, interviews, blog-like articles, news articles, conference proceedings, citable databases, etc.) or for journals not indexed on PubMed. Nevertheless, self-referencing rates (as well as the total number of publications by journal in Fig S1A) should only be viewed as estimates rather than exact figures. Despite this limitation, estimates of self-referencing rates from automated PubMed queries are quite consistent with values derived from manual calculation of self-referencing rates (Fig S1). Second, although the main focus has been on journals that predominantly publish experimental research, the full dataset of publications does include reviews, author corrections, and other article types. These alternative article types may exhibit distinct self-referencing rate distributions but are not distinguishable using the employed methodology. However, the effects of these alternative article types are mitigated by two main factors: 1) the criterium that publications required a minimum of 20 total references for inclusion in all analyses performed, which disproportionately eliminated publications such as author corrections and “letter to the editor” publications since they typically had few total references; and 2) failed PubMed queries, which disproportionately corresponded to non-research article/reference types. Third, although the individual PubMed query method improves author disambiguation compared to the use of databases with author name initials, author disambiguation is an ongoing challenge among bibliometric studies and may affect both self-referencing rates (potentially overestimating rates for authors with unusually common names) and matching of authors to the ranked-authors database. Authors could not be disambiguated by author affiliation due to inconsistent listing of author affiliations across PubMed publications. Fourth, comparison of self-referencing rates to total publications and fraction of publications self-referenced is imperfect, since publication dates were not accessible in the corresponding ranked-authors database. Ideally, for each publication with a calculated self-referencing rate, only the total publications prior to that publication would be included. However, the total number of career publications is considered a reasonable proxy for this purpose. Fifth, authors who frequently and/or selectively publish in the journals evaluated in this study are more strongly weighted than others. However, given the scope of this study (>94k publications evaluated), any single author is unlikely to have a large effect, particularly since most authors publish their work across a wide variety of journals. Finally, this study is clearly limited in scope to a small subset of journals. The data indicate that, while key indicators of self-referencing rates tend to fall within a relatively narrow range, there are also minor differences between journals which may reflect differences in publishing practices or content matter. These differences do not preclude the use of self-referencing guidelines, but rather indicate that thoughtful consideration (including aforementioned situational exceptions) should be exercised when authors, peer reviewers, and journals apply these statistical guidelines to individual cases. On the other hand, these data also reveal an advantage of studying self-referencing at the journal level: studies that combine publications across journals according to a field of study are inevitably combining sets of publications that were subject to distinct editorial regulations (e.g. article length, reference number, etc.), which have a demonstrable effect on self-referencing rates. Since publishers vary in the number of publications produced per year, publishers with high publishing rates will have the greatest weight in calculated self-referencing rates for field-grouped publication sets.

## Methods

### Data Acquisition and Processing

A complete list of PubMed IDs (PMIDs) was obtained from the NCBI website (https://ftp.ncbi.nlm.nih.gov/pub/pmc/) on 1/4/2023. For all PMIDs corresponding to the pre-defined set of journals, author lists were retrieved for each publication by a separate PubMed query. Subsequent queries were performed for all references within each publication to retrieve author lists for each reference. All PubMed queries were performed between 11/3/2022-1/23/2023 and included all publications for the entirety of 2022 for each journal. PubMed queries were performed using the Entrez module in the Biopython package [40]. Failed PubMed queries for both primary publications and referenced publications were repeated a minimum of 10 times.

For each successfully retrieved primary publication, a self-referencing rate was calculated for each author using a sequential string matching procedure. First, author names were checked for an exact string match. If no match was found, author names were subsequently truncated to various forms of first/middle initial and full surname and attempted to match against authors of the referenced publication (e.g. “John M. Smith” was truncated to “J M Smith”, “JM Smith”, and “J Smith”). Note that this subsequent step only identifies matches if the author names in the referenced publication are in the form of initials in the official PubMed query results. Although PubMed provides a downloadable open-access subset of full-text articles, the references therein only contain names with first/middle initials and full surname. Therefore, the method of performing individual PubMed queries was preferred over the more extensive downloadable database, as individual queries typically retrieve full author names which improve author disambiguation.

Most recent (2021) journal impact factors were obtained manually for each journal from https://jcr.clarivate.com/jcr/home, accessed on 1/22/2023.

The ranked-author database was downloaded from the external link in the corresponding publication ([33]; database version 5; last updated 10/28/2022). For all analyses involving author rank and/or total publications derived from the ranked-author database, only data corresponding to authors of the publications in this study’s dataset were included. Author rank was derived from the composite score corresponding to career-long, non-self citations only. Authors with a numerical rank >1 million were excluded from analyses, as author rank values become increasingly sparse.

### Calculation of Quartiles and Sliding Means

All percentile and central tendency values were calculated using the built-in Python “statistics” module or the “scoreatpercentile” module in the Scipy package with default parameters [41]. To estimate shifts in percent self-reference distributions as a function of a second variable (e.g. total references; Fig 2), quartile values (i.e. Q1, median, and Q3) were calculated for the distribution of percent self-references at each point along the *x*-axis (independent variable). Curves were then smoothed by calculating a sliding mean. Different window sizes (*W*) for sliding means were required to account for differences in the scale of the independent variable: *W*=11 for total references (Fig 2) and number of authors (Fig 3); *W*=21 for total publications (Fig 5A); *W*=5 for percent of all “self” publications referenced (Fig 5B) and percentage of authors appearing in the ranked-authors database (Fig 6B); and *W*=101 for author rank (Fig 5C). Quartile values for percent self-references were only calculated if 3 or more publications had the specified number of total references (x-axis). Linear interpolation was performed across gaps in the data, with the following maximum gap sizes (*G*): *G*=10 for total references (Fig 2) and number of authors (Fig 3); *G*=50 for total publications (Fig 5A); *G*=5 for percent of all “self” publications references (Fig 5B) and percentage of authors appearing in the ranked-authors database (Fig 6B); and *G*=5000 for author rank (Fig 5C). For all independent variables, linear interpolation was only necessary when reaching high x-axis values, where data become more sparse. When applicable, sliding means were calculated until a gap larger than the maximum gap size was encountered. For non-integer independent variables (e.g. percent of all “self” publications referenced), raw data were grouped into whole-integer bins using the numpy digitize function with *right=True* prior to calculating quartile values and sliding means.

## Data Availability

All code required to reproduce data in this publication are available at https://github.com/RossLabCSU/SelfReferencingRates2023. Additional data is provided as supplementary data accompanying this publication and at the provided Github repository.

## Supplementary Figure Legends

**Figure S1.**
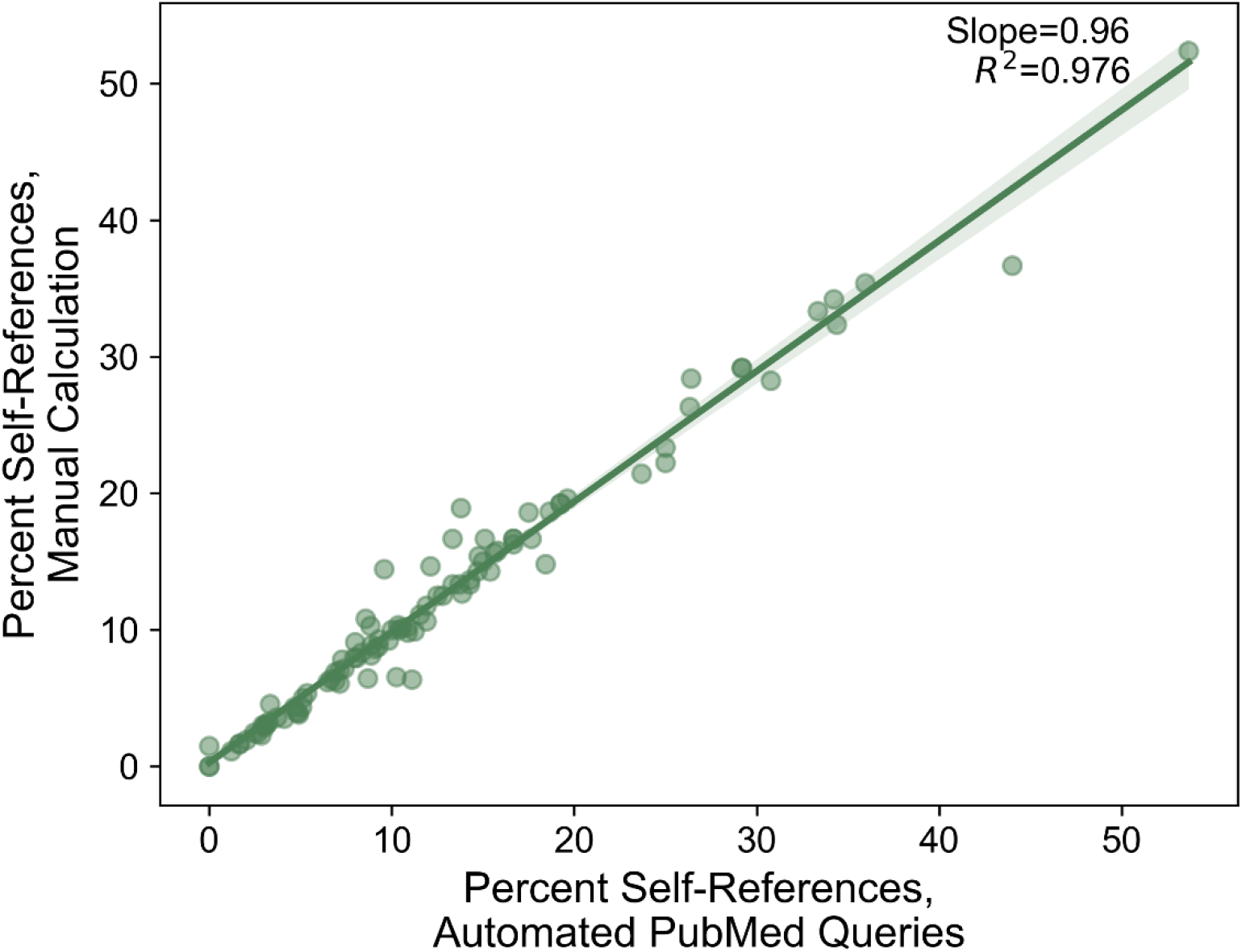
Comparison of manual versus automated calculation of self-referencing rates. A total of 100 publications were randomly selected from all publications evaluated. Self-referencing rates were calculated manually by examining the reference list in each publication, then compared to the self-referencing rate calculated via the automated PubMed query pipeline.

**Figure S2.**
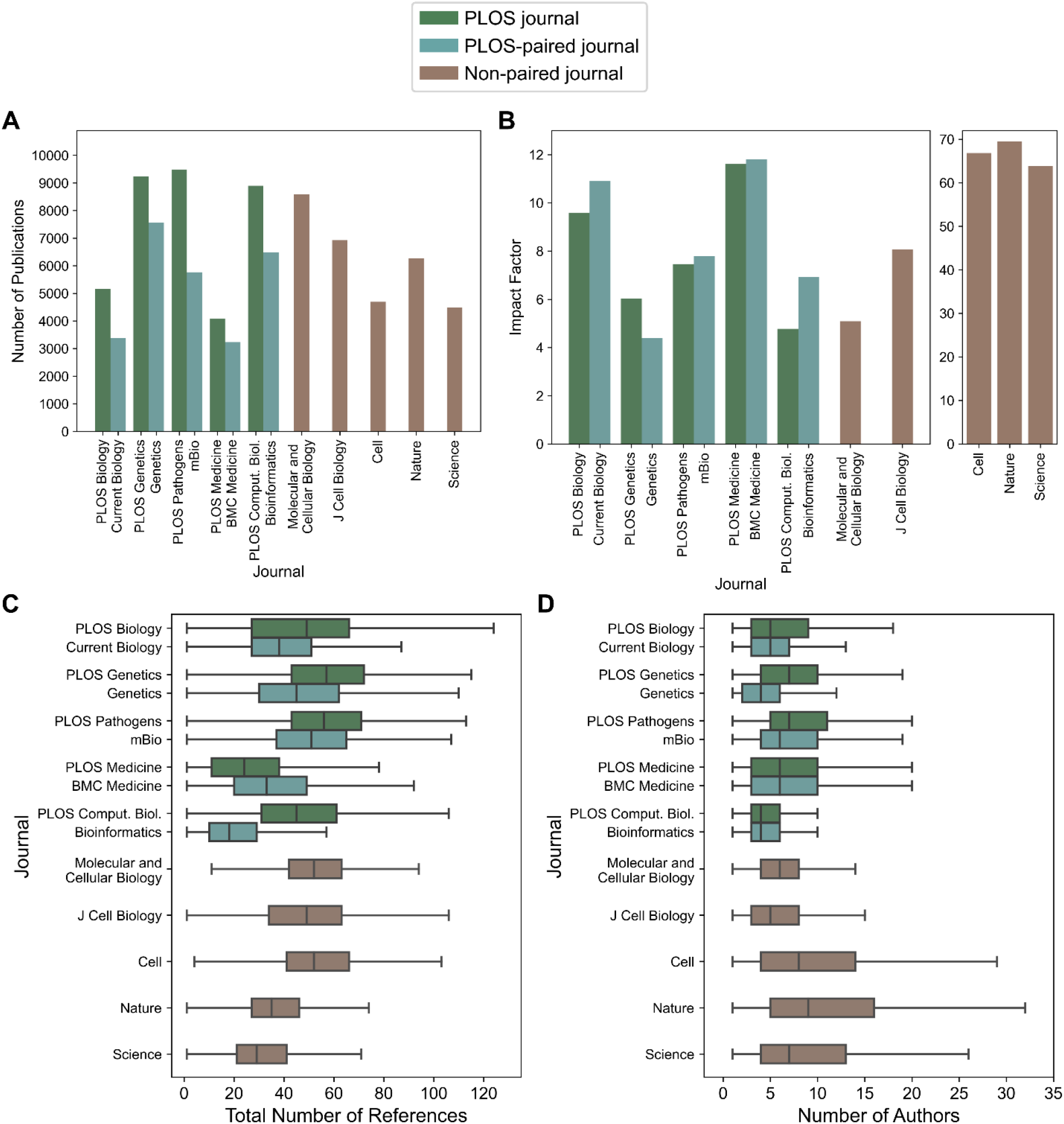
Comparison of number of publications, impact factor, number of references, and number of authors across journals. (A) Total number of publications evaluated in this study for each journal. (B) Most recent (2021) impact factor of each journal. (C) Distributions of total number of references within the primary publications evaluated for each journal. (D) Distributions of the number of authors for the primary publications evaluated for each journal.

**Figure S3.**
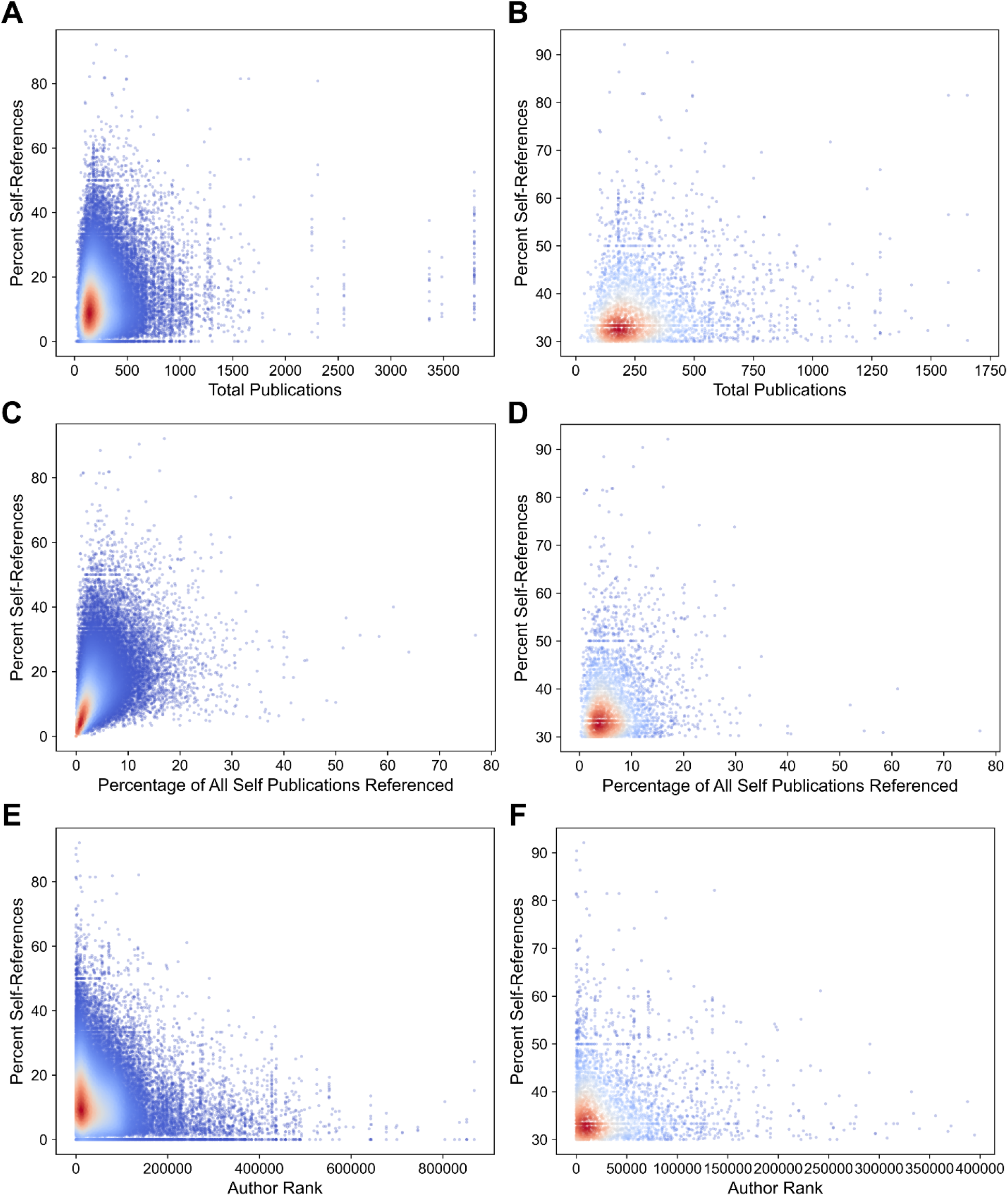
Self-referencing rates as a function of total publications, fraction of total publications referenced, and author rank. (A) Scatterplot depicting the percentage of self-references versus total publications. (B) Scatterplots focusing on the percentage of self-references versus total publications only for publications with very high self-referencing rates (≥30%) and authors with fewer than 2000 total publications. (C,D) Similar to panels A and B, but with the fraction of total publications self-referenced serving as the *x*-axis variable. (E,F) Similar to panels A and B, but with author rank [9,33] serving as the *x*-axis variable. For all panels, only the subset of authors appearing in the ranked-author database were included in analyses. Author rank in panels E and F was limited to 1 million and 400k, respectively, for visual clarity only. Point density was estimated using a kernel density estimate and colored accordingly, with dark red indicating high-density regions and dark blue indicating low-density regions.

**Figure S4.**
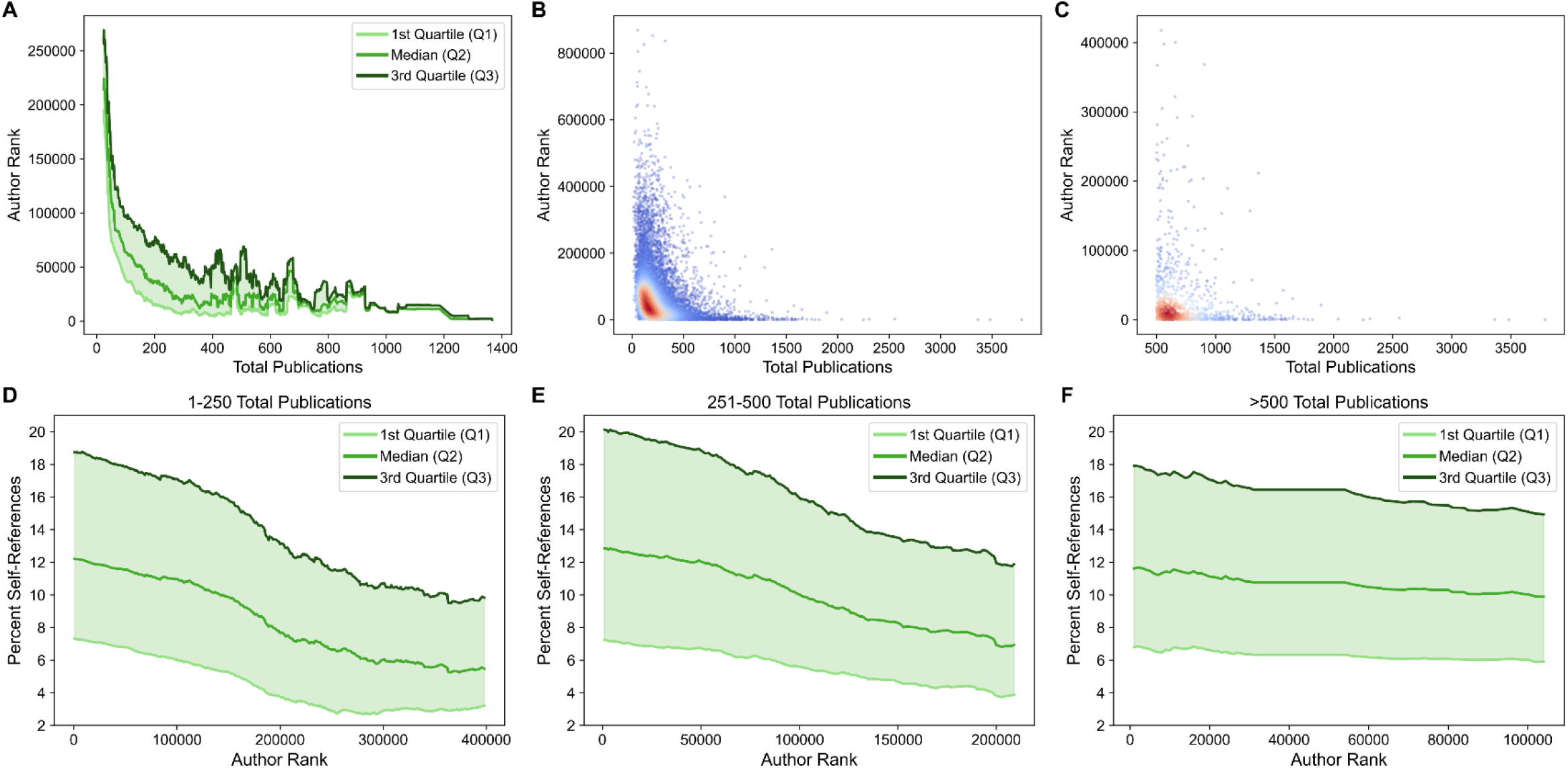
Author rank versus total publications. (A) Plot depicting shifts in author-rank quartile values as a function of total publications. (B) Scatterplot depicting author rank as a function of total publications. (C) Scatterplot of author rank versus total publications, focusing on authors with ≥500 total publications. Authors with an unusually large number of total publications are concentrated among top-ranking authors. For panels B and C, point density was estimated using a kernel density estimate and colored accordingly, with dark red indicating high-density regions and dark blue indicating low-density regions. (D-F) Shifts in percent self-reference distributions as a function of author rank, grouped by total publications as indicated in the title of each panel.

## Supplementary Table Legends

**Table S1. Estimated self-referencing rates for each publication.** Publications are listed by journal and PubMed ID. Authors are listed in the order that they appear in the original article, with each author separated by a semicolon. For each author, the number of self-references and percentage of self-references are estimated separately and appear in the same format as the listed authors (i.e. separated by semicolons, in the same order as the authors). Note that the number of self-references and the total number of references only reflect references that were successfully retrieved from PubMed queries: references not indexed on PubMed were excluded from analyses.

**Table S2. Comparison of automated versus manually calculated self-referencing estimates.** 100 publications were randomly selected from all publications analyzed in this study. Columns are as indicated for Table S1, with the addition of three new columns containing manually calculated values for the number of self-references, the total number of references in each publication, and the percentage of self-references.

**Table S3. Self-referencing rate summary by journal.** For each journal, the overall self-referencing rate at each indicated percentile is shown (columns 4-15). Additionally, the overall median self-referencing rate is indicated for each publication year from 2003-2022 (inclusive; columns 16-35).

**Table S4. Self-referencing rate values at percentiles as a function of total number of references.** For each journal, publications were binned based on the total number of references in each publication. Self-referencing rates were then calculated at each percentile (from 1-100) separately for each total-references bin. For columns 2-233, the column header indicates the total number of references, with values corresponding to the self-referencing rate estimated for the percentile indicated in column 2. The 25^th^, 50^th^, and 75^th^ percentile represent the Q1, median (Q2), and Q3, respectively, as shown in Fig 2. The 100^th^ percentile represents the maximum self-referencing rate. Self-referencing rates within each total-references column were smoothed using a sliding mean (see Methods). “nd” represents “no data”.

**Table S5. Self-referencing rate values at percentiles as a function of number of authors.** For each journal, publications were binned based on the number of authors on each publication. Self-referencing rates were then calculated at each percentile (from 1-100) separately for each number-of-authors bin. For columns 2-102, the column header indicates the number of authors, with values corresponding to the self-referencing rate estimated for the percentile indicated in column 2. The 25^th^, 50^th^, and 75^th^ percentile represent the Q1, median (Q2), and Q3, respectively, as shown in Fig 3. The 100^th^ percentile represents the maximum self-referencing rate. Self-referencing rates within each number-of-authors column were smoothed using a sliding mean (see Methods). “nd” represents “no data”.

**Table S6. Progressive contributions to self-referencing rates.** For each publication, progressive self-referencing rates – defined as the number or percentage of unique self-references contributed by each author, starting with the last author – were assigned to each author.

## Acknowledgements

I thank Dr. Eric D. Ross and Dr. Jennifer G. DeLuca for fruitful discussion, and Lindsey Bush for editing suggestions.

## Notes

### Competing Interest Statement

The authors have declared no competing interest.

